# Hypoxia preconditioned neural xenografts promote repair of brain tissue after stroke

**DOI:** 10.1101/2025.09.30.679547

**Authors:** Nora H. Rentsch, Rebecca Z. Weber, Beatriz Achón Buil, Chantal Bodenmann, Kathrin J. Zürcher, Vanessa Budny, Melanie Generali, Christian Tackenberg, Ruslan Rust

## Abstract

Stroke is a leading cause of long-term disability, yet no effective regenerative therapies exist. While cell-based therapies have shown promise in preclinical animal models, their clinical application remains limited due to poor survival of transplanted cells in the ischemic stroke environment. Hypoxic preconditioning has emerged as a strategy to potentially enhance graft survival, but the cellular mechanisms and translational relevance in human iPSC-derived neural progenitor cells (NPCs) are not fully understood. Here, we tested whether hypoxic preconditioning of NPCs affects their molecular and functional properties including proliferation and survival *in vitro* and after transplantation into a stroke mouse model. Hypoxic preconditioning enhanced proliferation and glial differentiation *in vitro*, improved cell survival post-transplantation, and enhanced regeneration-associated tissue responses such as vascular remodeling in the peri-infarct brain. These findings suggest that hypoxic preconditioning is a clinically translatable approach to increase the NPC graft survival in the post-stroke brain.

## INTRODUCTION

Stroke is a major cause of death and disability worldwide, yet current therapies including thrombolysis and mechanical thrombectomy, benefit only a fraction of patients because of a narrow treatment window^1,2^.

Stem cell therapy has emerged as a novel and promising paradigm to replace damaged neural circuitries after ischemic stroke^3^. Human induced pluripotent stem cell (iPSC)–derived neural progenitor cells (NPCs) offer an expandable, clinically compatible source for neural repair^4^, still fewer than 5% of grafted cells survive in the hostile post-ischemic niche^5,6^.

Hypoxic preconditioning (HP) has been proposed to enhance graft survival^7^ by activating hypoxia-inducible pathways that reduce apoptosis, increase angiogenesis, and modulate inflammation^8–10^. Short hypoxia (2–48 h) improves viability of several stem cell types *in vitro* and yields functional gains after transplantation^9–11^, yet its effects on human NPCs and the underlying mechanisms remain incompletely defined. In particular, long-term HP in NPCs has not been tested, transcriptomic changes that may influence metabolic pathways or glial versus neuronal fates are unknown, and the consequences for vascular remodeling are not established.

Here, we address these gaps by first exposing NPCs to either acute or sustained HP, profiling their metabolic, proliferative, and differentiation states *in vitro*. Next, we transplanted NPCs into a mouse model of photothrombotic stroke. We used longitudinal bioluminescence imaging with histology, transcriptomics and vascular quantification to determine whether HP enhances graft survival and modifies reparative tissue responses over a 4-week time course.

## RESULTS

### Generation of iPSC-derived NPCs and establishment of hypoxic preconditioning

First, we generated iPSC-derived NPCs under transgene-and xeno-free conditions and confirmed robust Nestin expression with no detectable Nanog expression, indicating a homogeneous progenitor population (**Fig. 1A,B**). NPCs were cultured either under atmospheric oxygen (20% O_2_) or in a low-oxygen gas mixture (1% O_2_, 5% CO_2_, 94% N_2_) to induce hypoxia (**Fig. 1C,D**). Live-cell hypoxia staining with Image-iT™ Green Hypoxia Reagent revealed strong fluorescence exclusively in low-oxygen cultures, confirming establishment of a stable hypoxic environment (**Fig. 1E,F**).

**Fig. 1:**
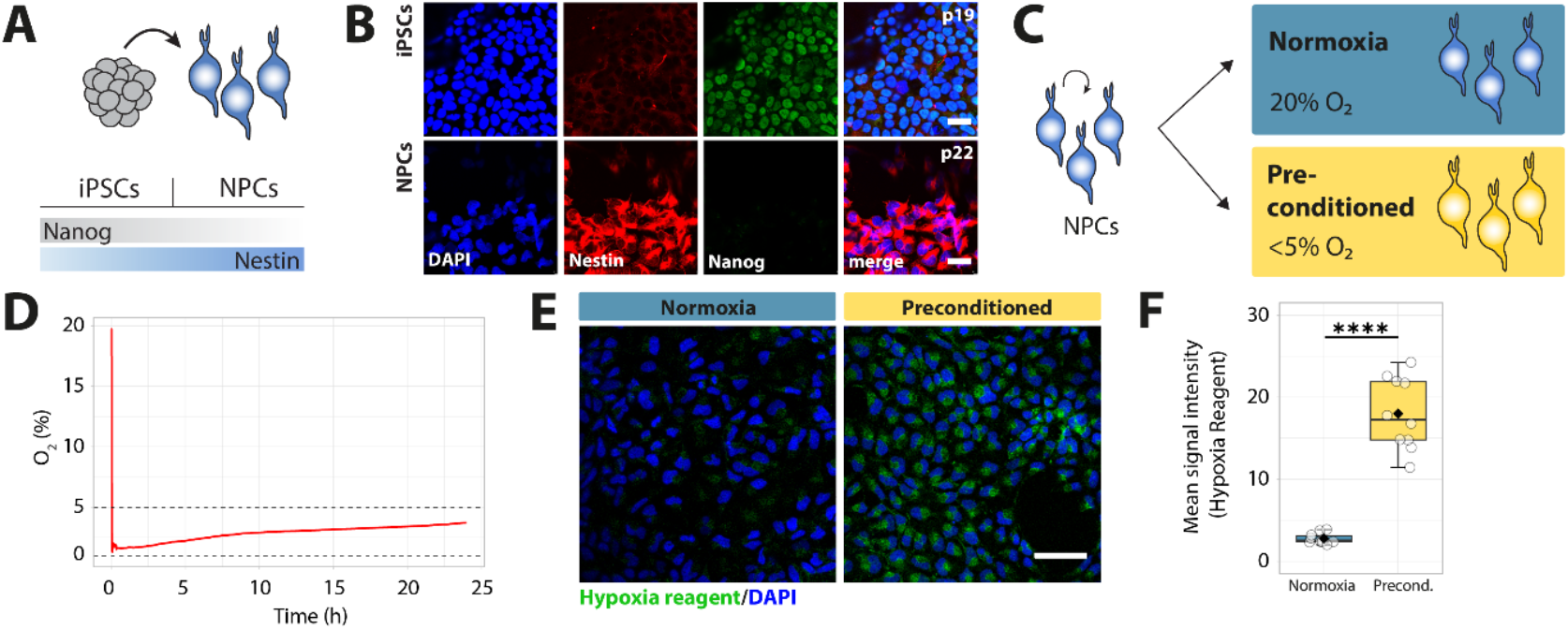
Validation of hypoxic environment in experimental settings. (**A**) Schematic of iPSC to NPC differentiation with representative marker expression profile. (**B**) Immunofluorescence confirming Nestin (red) expression in NPCs and Nanog (green) expression in iPSCs. Scale bar, 20 µm. (**C**) Experimental design for normoxic (20% O_2_) and hypoxic preconditioning (<5% O_2_) of NPCs. (**D**) Oxygen concentration profile in the hypoxic chamber over 24 h after flushing with low oxygen gas mixture. (**E**) Live-cell staining with Image-iT™ Green Hypoxia Reagent (green) and DAPI (blue) showing hypoxia-specific signal in preconditioned cultures. Scale bar, 50 µm. (**F**) Quantification of hypoxia reagent fluorescence intensity in normoxic and preconditioned NPCs after 4 days. In F: Differences between normoxic and hypoxic cells were assessed using an unpaired two-sided t-test. Boxplots show median (center line), interquartile range (box), and minimum-to-maximum values (whiskers); each point represents one analyzed image. ****p<0.0001.

### Hypoxic preconditioning increases cell proliferation and drives NPC towards a glial fate *in vitro*

To understand how cells react to a hypoxic environment, NPCs were exposed to low oxygen for (a) 14 days during the final stage of neuronal induction (defined here as acute hypoxia, AH-NPCs) or (b) 4 days during proliferation and, subsequently, 14 days during neuronal induction (defined here as hypoxic preconditioning, HP-NPCs), similar to established preconditioning paradigms.^12^ A 4-day hypoxic period during proliferation is sufficient to activate supporting pathways to promote viability and manifest effects over several cell cycles before switching to neural induction. Since early neural markers are expressed within 1-2 weeks of differentiation, a 14-day induction window is reasonable to capture the impact of hypoxia on neural stem cell fate. As a control, NPCs were cultured under atmospheric conditions (defined here as normoxia, N-NPCs, **Fig. 2A**).

**Figure 2.**
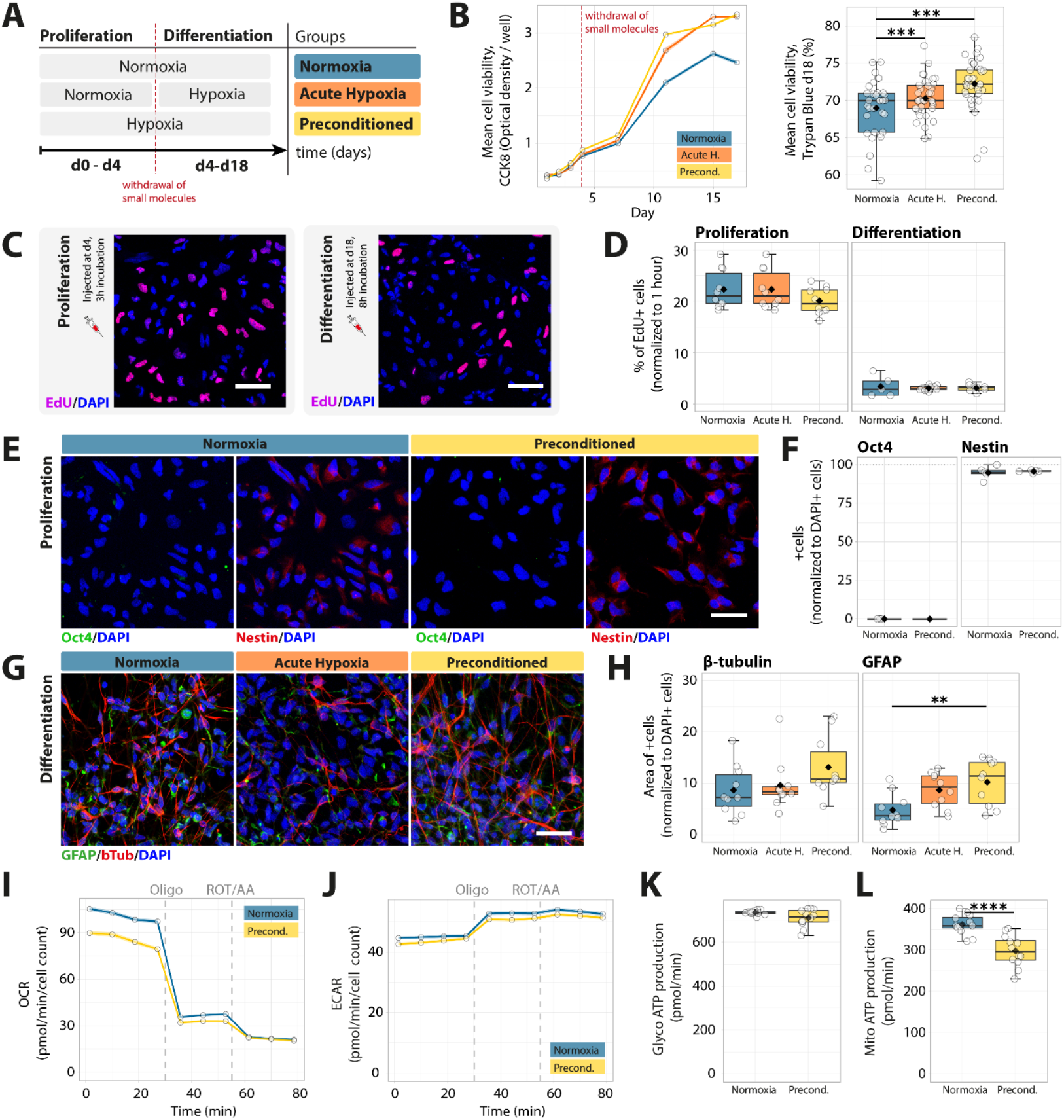
Hypoxic preconditioning increases cell viability and biases NPC fate towards glia *in vitro*. (**A**) Experimental design showing normoxic (20% O_2_), acute hypoxia (<5% O_2_ during differentiation only), and hypoxic preconditioning (<5% O_2_ during proliferation and differentiation) groups. (**B**) Left: Mean cell viability (optical density) during culture measured with Cell Counting Kit 8 – assay (CCK8); day 4 marks the induction of differentiation (dashed red line). Right: Cell viability measured with Trypan Blue assay at day 18. (**C**) Representative EdU incorporation (magenta) in proliferative and differentiated NPCs. Scale bar, 50 µm. (**D**) Quantification of EdU+ cells during proliferation and differentiation phases. (**E**) Immunofluorescence for Oct4 (green) and Nestin (red) in proliferating NPCs. Scale bar, 30 µm. (**F**) Quantification of Oct4+ and Nestin+ cells. (**G**) Representative images of differentiated NPCs stained for βIII-Tubulin (red) and GFAP (green). Scale bar, 30 µm. (**H**) Quantification of βIII-Tubulin+ and GFAP+ cells. (**I**,**J**) Oxygen consumption rate (OCR) and proton efflux rate (PER) measured by Seahorse XF24 analyzer; oligomycin (Oligo) and rotenone/antimycin A (ROT/AA) added at indicated times. (**K**,**L**) Quantification of glycolytic (glycoATP) and mitochondrial (mitoATP) ATP production. In F, K, L: Differences between control and hypoxic cells were assessed using an unpaired two-sided t-test. In B,D,H: Differences between N-NPCs, AH-NPCs, HP-NPCs were assessed using ANOVA with tukey HSD post-hoc test. In all boxplots, center line = median, box = interquartile range, whiskers = minimum–maximum, and each point represents one analyzed FOV (D, F, H) or one analyzed well (K, L). *p<0.05, **p<0.01, ***p<0.001, ****p<0.0001.

A metabolic assay (Cell counting kit-8, CCK8) was conducted during proliferation and differentiation to longitudinally examine cell viability. Whereas similar viability between groups was observed during proliferation (day 1 to 4), viability increased in HP-NPCs and AH-NPCs compared to N-NPCs 3 days after inducing differentiation (day 7: p<0.05, day 11/ 15 /17: p< 0.0001). These results were supported by an independent non-metabolic Trypan Blue viability assay at day 18 that showed increased viability in hypoxic treated NPCs (AH-NPCs: p = 3*10^-11^ and HP-NPCs: p = 7.9*10^-12^) compared to N-NPCs (**Fig. 2B**). Overall, this data indicates an increased cell viability under a hypoxic environment.

Cells were treated with 5-ethynyl-2’-deoxyuridine (EdU), a nucleotide analog that incorporates into dividing cells, to assess their proliferation ratio. As expected, proliferation was significantly higher when cells were cultured under proliferative conditions (22% of EdU-positive cells) compared to after two weeks of differentiation (3% of EdU-positive cells) across all groups (N-NPC: p=1.23*10^-7^, AH-NPC: p=4.36*10^-12^, HP-NPC: p=1.12*10^-13^). No significant differences were observed between the treatment groups within either proliferation or differentiation phase, indicating that hypoxic exposure did not alter proliferation rates under these culture conditions (**Fig. 2C,D**).

Next, we phenotyped cells under proliferation and differentiation conditions. Immunocytochemistry revealed strong Nestin expression (>90%) and complete absence of Oct4 under proliferative conditions across all groups (**Fig. 2E,F**). Under differentiation conditions, the proportion of GFAP-positive cells increased with hypoxia (N-NPC = 4.8%, AH-NPC = 8.7%, HP-NPC = 10.2%), with a significant HP-NPC to N-NPC increase (p = 0.0078), suggesting a shift towards a glial fate (**Fig. 2G,H**). However, βIII-Tubulin-positive neurons remained more abundant than astrocytes in all groups, and Olig2 staining did not reveal oligodendrocyte generation (**Supp. Fig. 1A**).

To assess whether a hypoxic environment induces changes in the energy metabolism of NPCs we measured the oxygen consumption rate (OCR) and extracellular acidification rate (ECAR) with a Seahorse XF24 extracellular flux analyzer (**Fig. 2 I,J**). Mitochondrial (mito) and glycolytic (glyco) ATP production were analyzed by first inhibiting complex V of the respiratory chain with oligomycin, followed by the inhibition of complexes I and III using rotenone and antimycin. Whereas there was no significant difference observed in the glycoATP production hypoxic preconditioning caused lower mitoATP production in NPCs compared to normoxic conditions (p<0.0001) (**Fig. 2 K,L**).

### Hypoxic preconditioning increases expression of regenerative factors in NPCs

Bulk RNA-seq was performed to dissect transcriptional mechanisms underlying hypoxia-induced changes in NPCs. We analyzed the transcript of NPCs across three conditions: N-NPC, AH-NPC, HP-NPC, during both proliferation and differentiation. Unsupervised heatmap clustering and PCA revealed that hypoxia induced clear transcriptomic divergence, particularly between HP-NPC and N-NPC (**Figure 3A-D**).

**Figure 3:**
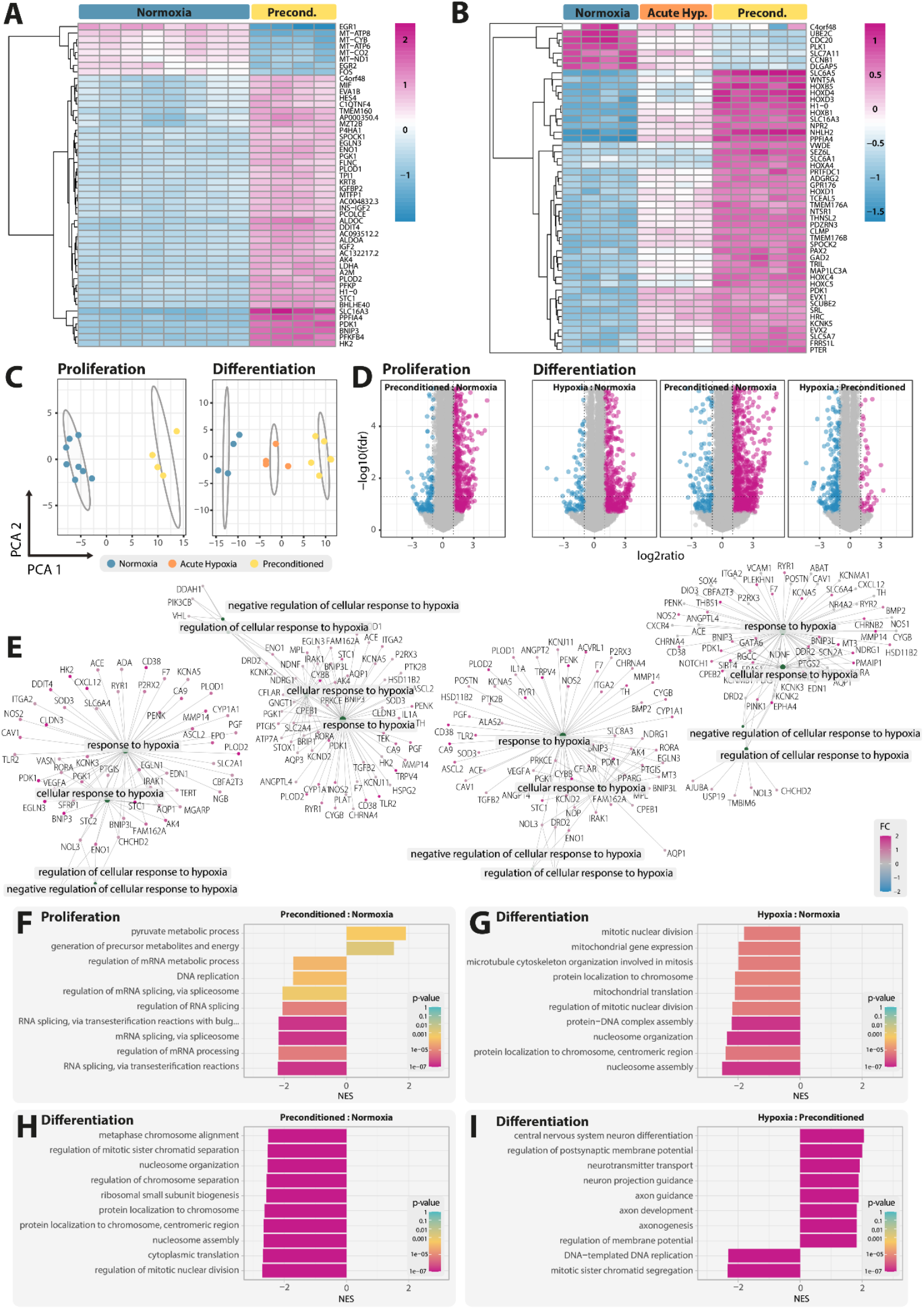
Hypoxic preconditioning alters transcriptional programs during NPC proliferation and differentiation. **(A–B)** Heatmaps showing differentially expressed genes (DEGs) during proliferation **(A)** and differentiation **(B)** of NPCs under normoxic (N), acute hypoxic (AH), or hypoxic preconditioning (HP) conditions (n = 4 - 8 per group). **(C)** Principal component analysis (PCA) reveals condition-specific transcriptomic clustering during proliferation and differentiation. **(D)** Volcano plots of DEGs comparing N vs. HP and N vs. AH during proliferation and differentiation, highlighting upregulated (magenta) and downregulated (blue) genes. **(E)** Gene regulatory network (GRN) analysis to identify hypoxia-responsive pathways **(F-I)** Gene set enrichment analysis (GSEA) indicates hypoxia-dependent regulation of key biological processes. p-value cut-off = 0.2.

Differential gene expression (DGE) analysis revealed pronounced transcriptional remodeling under hypoxic conditions. During proliferation, hypoxic preconditioning altered 793 transcripts (705 up, including glycolytic and hypoxia-adaptive regulators SLC16A3, PDK1, BNIP3; 88 down, including neurodevelopmental genes PAX3, NRG1). In differentiated cultures, acute hypoxia modulated 458 genes (388 up, including synapse-and signaling-associated PPFIA4, NPR2, chromatin regulator H1-0; 70 down, including cell cycle genes PLK1, PIF1), while hypoxic preconditioning elicited the most extensive response, altering 1,340 genes (1,075 up, including neurogenesis-and ECM-related PDZRN3, PPFIA4, SPOCK2; 265 down, including mitotic regulators BUB1, KIF20A). In contrast, direct comparison between HP-NPC and AH-NPC during differentiation revealed only 313 changes (52 up, including Wnt and transcriptional regulators SFRP2, IKZF1; 261 down, including synaptic and neurotrophin receptors SEZ6L, NTRK3), indicating a possible convergence of transcriptional programs once cells adapt to sustained low oxygen exposure.

Gene set enrichment analysis (GSEA) first confirmed major upregulation of networks associated with hypoxia during proliferation and differentiation (**Fig 3E**). Apart from hypoxia-related pathways, we identified marked shifts in biological processes during proliferation and differentiation. During proliferation, HP-NPCs upregulated pathways related to pyruvate metabolic process, generation of precursor metabolites, while downregulating cell cycle and DNA replication programs compared to N-NPCs (**Fig. 3F**). In differentiated cultures, hypoxia (AH-NPC and HP-NPC) downregulated pathways related to mitotic and chromosomal organization pathways compared to normoxia (**Fig. 3G-H**).

In contrast, comparing hypoxia (AH-NPC) to preconditioning (HP-NPC) in the differentiated state revealed HP-NPC-specific enrichment of neuronal maturation and synaptic pathways, including central nervous system neuron differentiation, neurotransmitter transport, axonogenesis, and regulation of membrane potential, indicating potentially upregulated differentiation program (**Fig. 3I**).

### Hypoxic preconditioning increases cell survival after transplantation into the ischemic brain

We designed four experimental cohorts of adult immunodeficient Rag2^-/-^ mice (**Fig. 4A**): the no stroke group received intracerebral injections of iPSC-derived NPCs expanded under normoxic conditions without photothrombotic stroke induction; the unconditioned and preconditioned groups both sustained a photothrombotic stroke followed by NPC transplantation, with cells expanded under normoxic (20% O_2_) or hypoxic (>5 % O_2_) conditions, respectively; and a no-cells group received a stroke but no cell therapy.

**Figure 4:**
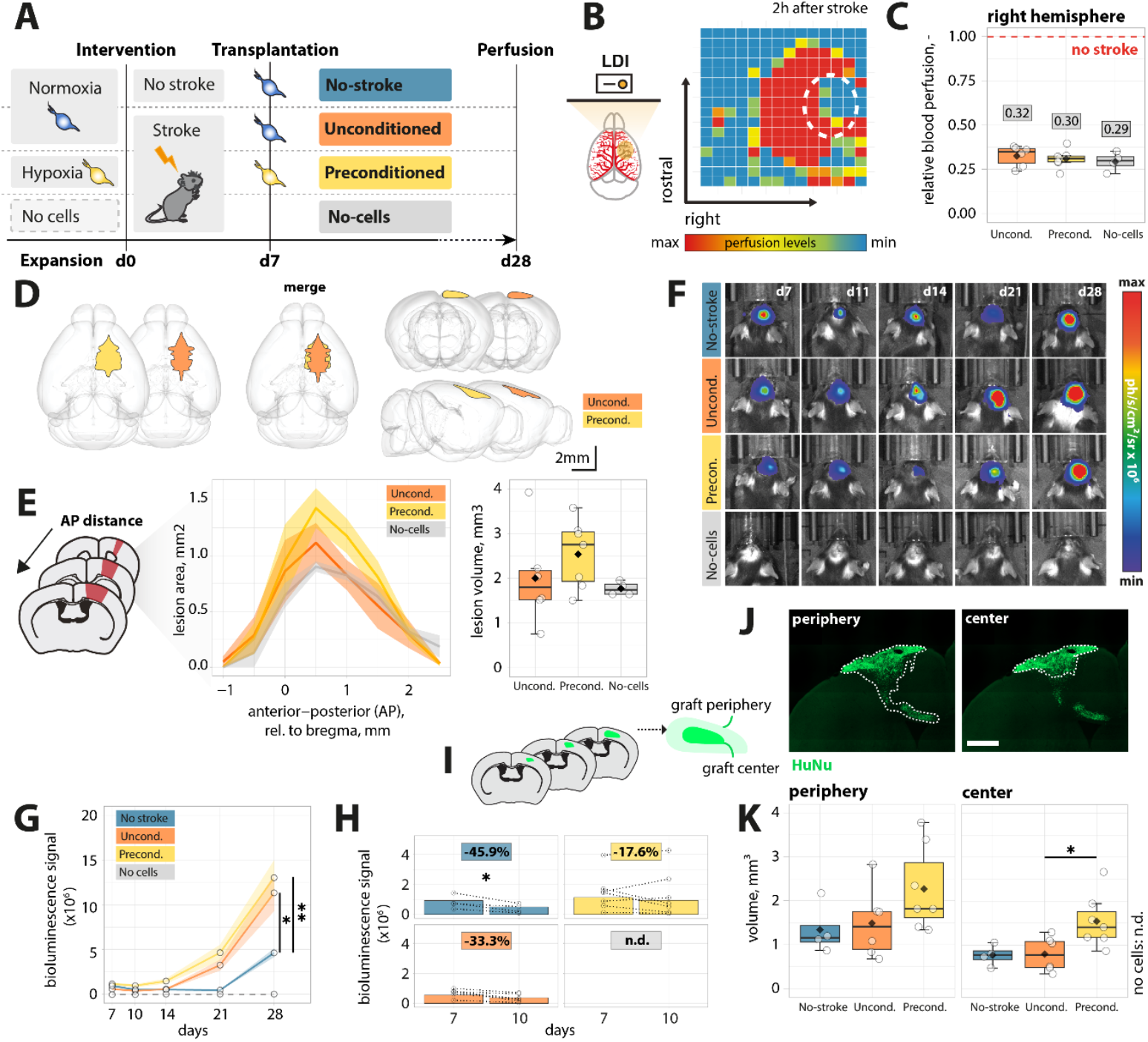
Hypoxic preconditioning increases NPC survival and alters graft distribution after transplantation into the ischemic brain. **(A)** Experimental design showing four groups: “No-stroke” (normoxic NPCs transplanted without stroke), “Unconditioned” (normoxic NPCs transplanted after stroke), “Preconditioned” (hypoxic NPCs transplanted after stroke), and “No-cells” (stroke without cell therapy). NPCs were expanded under normoxia (20% O_2_) or hypoxia (<5% O_2_). Photothrombotic stroke was induced at day 0, intracerebral transplantation performed at day 7, and perfusion at day 28. **(B)** Representative Laser Doppler imaging (LDI) heatmap of cerebral blood flow 2 h after stroke. **(C)** Quantification of relative cerebral perfusion in the right hemisphere. **(D)** Schematic of coronal section series used for stroke volume analysis. **(E)** Rostrocaudal lesion profiles and quantification of stroke volume (mm^3^). Differences between unconditioned, preconditioned and control subjects were assessed using ANOVA with tukey HSD post-hoc test. **(F)** Representative bioluminescence imaging (BLI) of grafted NPCs at d7, 10, 14, 21 and 28 post-stroke. Color scale represents radiance (photons s^−1^ cm^−2^ sr^−1^). **(G)** Longitudinal quantification of BLI signal over 21 days. **(H)** Change in BLI signal between day 7 and 10 post-injury. **(I)** Schematic defining “graft core” (≈90% of HuNu+ cells) and “graft periphery” (remaining 10%). **(J)** Representative HuNu immunofluorescence images showing graft periphery and core. Scale bar, 500 µm. **(K)** Quantification of graft periphery and core volumes (mm^3^) at day 28. In C: Differences between unconditioned, preconditioned and no-cell control subjects were assessed with paired Wilcoxon-Mann-Whitney test. In K: Differences between unconditioned, preconditioned and unstroked control subjects were assessed using ANOVA with tukey HSD post-hoc test. In G: Differences between unconditioned, preconditioned and unstroked control subjects were assessed using ANOVA with estimated marginal means (EMM) post-hoc test and bonferroni correction. In H: Differences between day 7 and 10 within groups were assessed using an unpaired two-sided t-test. In all boxplots, center line = median, box = interquartile range, whiskers = minimum–maximum, and each point represents one animal. *p<0.05, *p<0.01.

To confirm equivalent ischemic severity at onset, we performed Laser Doppler imaging immediately after stroke induction (**Fig. 4B**). Quantitative analysis (**Fig. 4C**) revealed a comparable reduction in relative cerebral perfusion in the right hemisphere across all stroked groups: 68±4% in the unconditioned cohort, 70±5% in the preconditioned cohort, and 71±3% in the no-cells cohort (mean ± SD; p>0.05). This confirms that initial ischemic burden did not differ between treatment groups.

Next, we assessed lesion expansion 28 days after injury, which corresponds to 21 days after cell transplantation, by measuring infarct area on serial coronal sections spanning-1 to +2.5 mm relative to bregma (**Fig. 4D**). Stroke volume was calculated by integrating the area across the rostrocaudal axis and reached 2.54±0.75 mm^3^ in the unconditioned group, 2.00±1.10 mm^3^ in the preconditioned group, and 1.77±0.15 mm^3^ in the no-cells group (mean ± SD; p>0.05) (**Fig. 4E)**. Thus, neither hypoxic preconditioning of NPCs nor cell delivery per se altered the extent of initial tissue damage.

To track viability of grafted cells longitudinally *in vivo*, NPCs were transduced with a dual red firefly luciferase (rFluc) – eGFP reporter. Immediately following grafting, comparable bioluminescent signals across groups confirmed equivalent cell delivery (**Fig. 4F**). Serial bioluminescent imaging (BLI) over 21 days demonstrated sustained survival of transplanted NPCs in all stroked animals (**Fig. 4G**). Focusing on the acute engraftment window (days 7 to 10, **Fig. 4H**), unconditioned NPCs exhibited a 33.3% decrease in luciferase signal, whereas preconditioned NPCs declined by only 17.6%. Noticeably, 3 out of 7 animals that received HP-NPCs showed increased luciferase signal within the early time window, suggesting enhanced early viability conferred by hypoxic preconditioning. In contrast, N-NPCs delivered into non-stroked brains showed a loss in signal of 45.9% (p = 0.0381) over the same period, with decreased signal observed in all animals, implying that the cells require the space in tissue induced by the stroke to expand efficiently.

21 days after cell delivery, surviving graft cells were identified using human-specific antibodies in coronal brain sections (**Fig. 4I**). Graft size was analyzed to investigate homing and potential migration processes of transplanted NPCs. On that account, the graft was quantified in its two areal components: the graft core, defined as the region where approximately 90% of grafted cells reside, and the graft periphery, in which the additional 10% were included (**Fig. 4J**). In general, transplanted NPCs survived and repopulated the lesion cavity within the 21-day time course, regardless of their culture regime. Our analysis revealed that the volume of the core component of grafted HP-NPCs was significantly increased compared to grafted unconditioned cells (Preconditioned: 1.54 mm^3^±0.6; unconditioned: 0.8 mm^3^±0.39, no stroke: 0.77 mm^3^±0.24, p=0.038, **Fig. 4K**). Lastly, we observed a broader dispersion of transplanted cells away from the injection site in the preconditioned group, possibly indicating increased migration (Preconditioned: 2.27 mm^3^±0.96; unconditioned:1.44 mm^3^±0.81, no stroke: 0.77 mm^3^±0.24, **Fig. 4K**).

### Hypoxic preconditioning enhances differentiation and synaptic integration of neural progenitor cell grafts after stroke

Immunohistochemistry was performed on coronal brain sections at 21 days post-transplantation to characterize the composition of the graft (**Fig. 5A**). Co-staining with antihuman nuclear antigen (HuNu) and cell lineage markers revealed no residual pluripotent cells: Nanog^+^/HuNu^+^ profiles were undetectable in any group (**Fig. 5B**). A small fraction of grafted cells retained progenitor identity, with Pax6^+^/HuNu^+^ double positive cells comprising 3.2 ± 0.9% of the preconditioned graft and 4.4 ± 1.1% of the unconditioned graft in stroked brains; the no-stroke group showed 3.1 ± 0.8% (mean ± SD; **Fig. 5C**). This indicates that hypoxic preconditioning neither increases residual pluripotency nor alter the spontaneous differentiation of progenitor cells.

**Figure 5:**
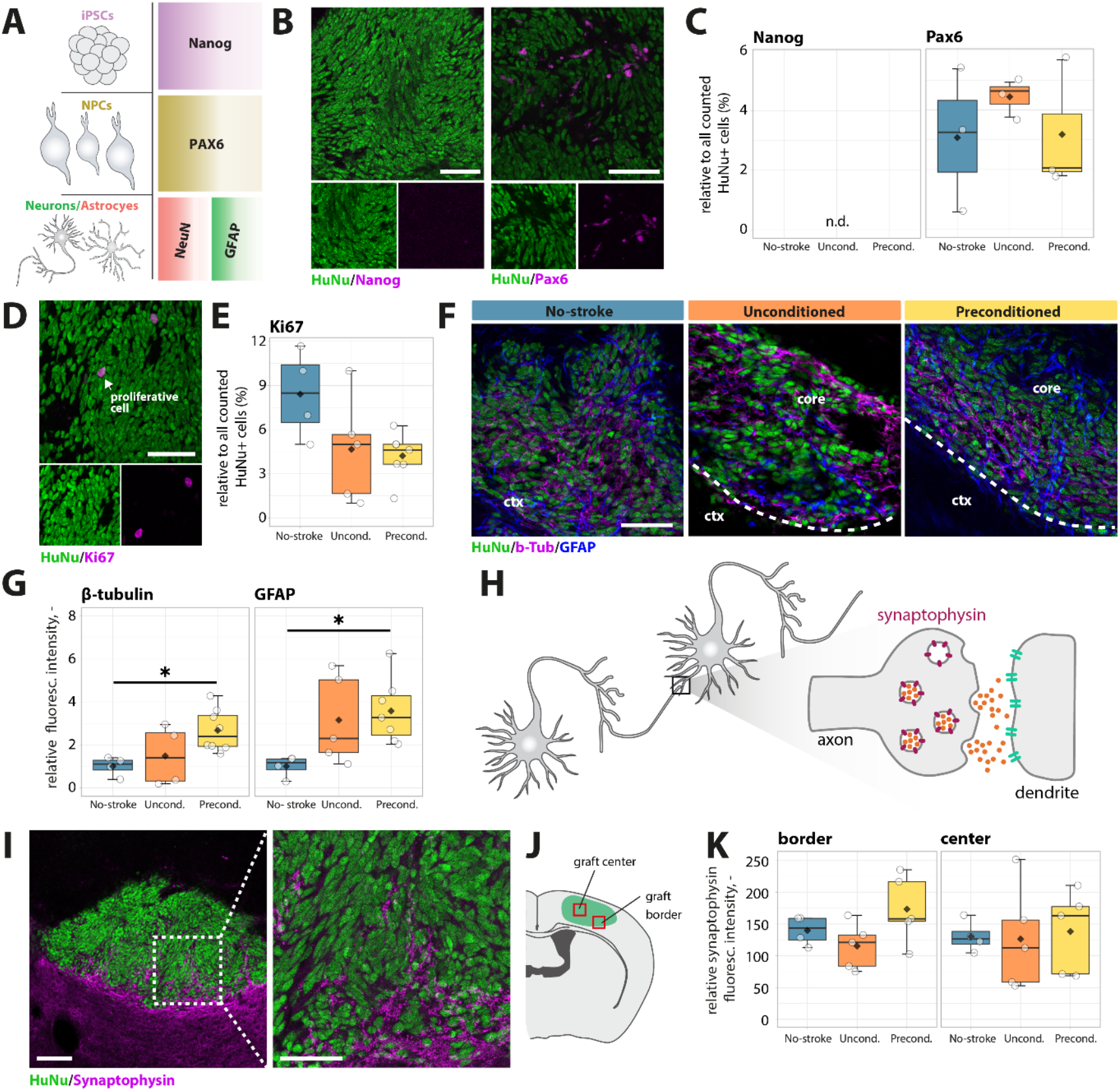
Hypoxic preconditioning influences graft composition in ischemic brain. (**A**) Schematic representation of analyzed cell types and protein markers used for characterization. (**B**) Immunofluorescence for Nanog and Pax6 (magenta) in graft, counterstained with HuNu (green). n.d. = not detected. Scale bar, 50 μm. (**C**) Quantification of Nanog^+^ and Pax6^+^ cells in graft. (**D**) Representative images of proliferating cells in graft stained with Ki67 (magenta). Scale bar, 50 μm. (**E**) Quantification of Ki67^+^ cells. (**F**) Fluorescence images of βIII-Tubulin and GFAP in graft core. Scale bar, 50 μm.(**G**) Quantification of fluorescence intensity of βIII-Tubulin and GFAP, normalized to the no-stroke group. (**H**) Schematic illustration of expression of synaptophysin in neuronal synapse. (**I**) Representative image of graft stained with Synaptophysin (magenta) and HuNu (green). Scale bar, 150 μm. The rectangle shows an enlarged section as displayed on the right. Scale bar, 50 μm.(**J**) Schematic of center and border in the graft. (**K**) Quantification of synaptophysin fluorescence intensity in the border (left) and center (right) of the graft. In C, E, G, K: Differences between unconditioned, preconditioned and no-stroke control subjects were assessed using ANOVA with tukey HSD post-hoc test. In all boxplots, center line = median, box = interquartile range, whiskers = minimum–maximum, and each point represents one animal. *p<0.05.

To assess graft cell proliferation as an indicator of potential tumorigenicity, Ki67 immunostaining was performed. Proliferation within the graft was similarly low: Ki67^+^/HuNu^+^ cells represented 4.1 ± 0.7% of preconditioned and 4.7 ± 0.6% of unconditioned grafts in stroked animals, whereas the no-stroke graft exhibited a higher proliferative index of 8.4 ± 1.2% (**Fig. 5D,E**). Only occasional Ki67^+^ cells lacking HuNu were observed in the host parenchyma. These findings indicate a low proliferation rate and thus minimal risk of tumor formation.

In stroked animals, preconditioned NPCs showed significantly greater relative fluorescence intensity for both βIIITubulin-(1.35 ± 0.15-fold vs unconditioned; p < 0.05) and GFAP (1.28 ± 0.12fold vs unconditioned; p < 0.05), whereas-no difference was found between unconditioned and preconditioned grafts (**Fig. 5F,G**). Finally, synaptic integration was evaluated by Synaptophysin/HuNu co--labeling at the graft border and center. At the graft– host interface, preconditioned cells exhibited a higher Synaptophysin signal (191 ± 40) compared to unconditioned cells (113 ± 41; p < 0.05), with the no-stroke group at an intermediate level (159 ± 0.1). In the graft core, Synaptophysin intensities were similar across preconditioned (144 ± 63), unconditioned (132 ± 83), and no--stroke-(134 ± 42) groups (**Fig. 5H–K**). Collectively, these data indicate that hypoxic preconditioning enhances neuronal and astrocytic differentiation and promotes synaptic integration at the graft border.

### Hypoxic preconditioning of cell grafts is associated with reduced inflammation

To identify stroke-related inflammation 28 days after injury, we analyzed level of the microglial marker Iba1 in three regions: in the stroke core, in the peri-infarct region (ischemic border zone, ibz) adjacent to the core and the contralesional cortex. Iba1 is upregulated upon activation of microglia and/or peripheral macrophages and plays a role during the proinflammatory response to ischemia and tissue injury. Immunostainings revealed an overall increased expression of Iba1 in the ipsilateral cortex in the core and the peri-infarct regions (**Fig. 6A-B**). A significant reduction of Iba1-expression was found in the stroke core of mice that had received preconditioned NPCs compared to the group that did not receive any cells (2.3-fold relative to control, p<0.01). As expected, low Iba1 expression was observed in the contralesional hemisphere, and no significant difference could be observed across all groups (p>0.05). These results suggest that hypoxic preconditioning of the transplanted cells attenuates inflammation in the tissue.

**Figure 6:**
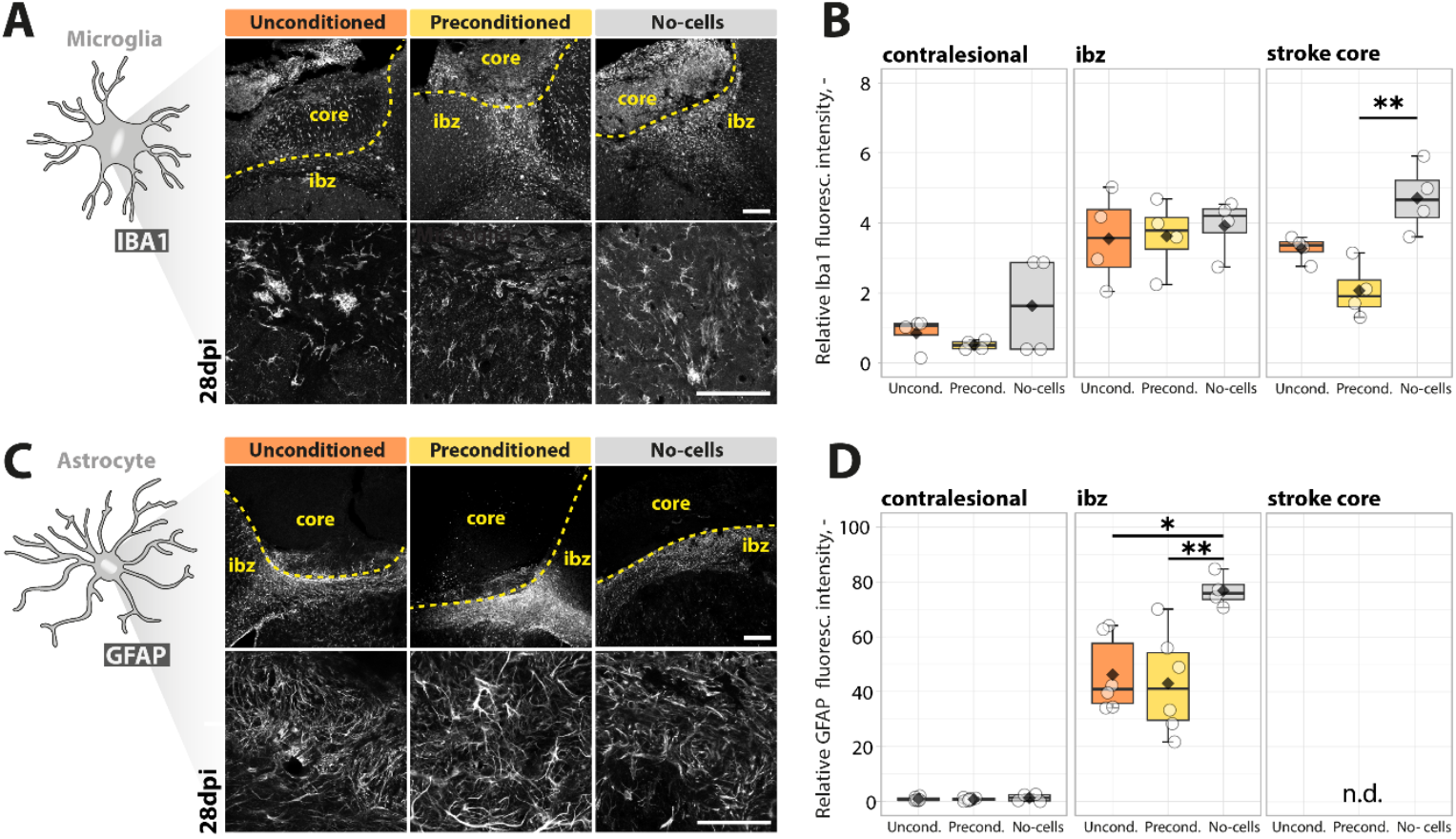
Tissue response is modified by hypoxic preconditioned cells. (**A**) Fluorescent images of Iba1 in stroke core and ischemic border zone (ibz) 28 days post-injury (dpi). The dotted line marks the border between the core and the ibz. Scale bar, 100 μm. The lower panel shows enlarged sections of the ibz. Scale bars, 100 μm. (**B**) Quantification of Iba1 fluorescence intensity in the contralesional site (left), ibz (middle) and stroke core (right). (**C**) Immunofluorescence for GFAP marker in tissue sections 28 dpi. Magnified images are displayed in the lower panel. Scale bars, 100 μm (**D**). Quantification of GFAP signal intensity. n.d. = not detected. In B, D: Differences between unconditioned, preconditioned and no-cells control subjects were assessed using ANOVA with tukey HSD post-hoc test. In all boxplots, center line = median, box = interquartile range, whiskers = minimum–maximum, and each point represents one animal. *p<0.05, **p<0.01.

Next, we quantified the fluorescence intensities of astrocyte marker GFAP in in the same regions (contralesional, ibz and stroke core) to identify the extent of stroke-related scarring (**Fig. 6C**). High GFAP expression levels were observed along the ischemic border zone (ibz) but the signal was reduced in the unconditioned group (p > 0.05) and in preconditioned group (p<0.01) compared to the no-cells group whereas no signal was detected in the core. (**Fig. 6D**). Negligible expression was observed in the contralateral hemisphere in all groups. These results are indicating glial scar formation and hence regenerative properties in the injured tissue.

### Hypoxic preconditioning of neural progenitor cell grafts enhances vascular repair after stroke

Next, we investigated several vascular parameters such as blood vessel distribution, blood vessel density, and blood vessel integrity (**Fig. 7A,B**). The overall vascular density, number of branches and total length of the vascular network was increased in the peri-infarct areas in mice that received NPCs compared to the control group (**Fig. 7C, D**). Most prominent, mice receiving preconditioned cells had an increase of +100% (p < 0.01) in vascular network density, +122% (p < 0.01) in the number of branches and +140% (p < 0.001) in the vascular length compared to the control group (no-stroke). Interestingly, vascular length was also significantly higher in the preconditioned group when compared to animals receiving unconditioned NPCs (+29%, p < 0.05). Importantly, no noticeable vascular differences between the groups were observed in the contralesional hemisphere (all p >0.05, **Supp. Fig 2**).

**Figure 7:**
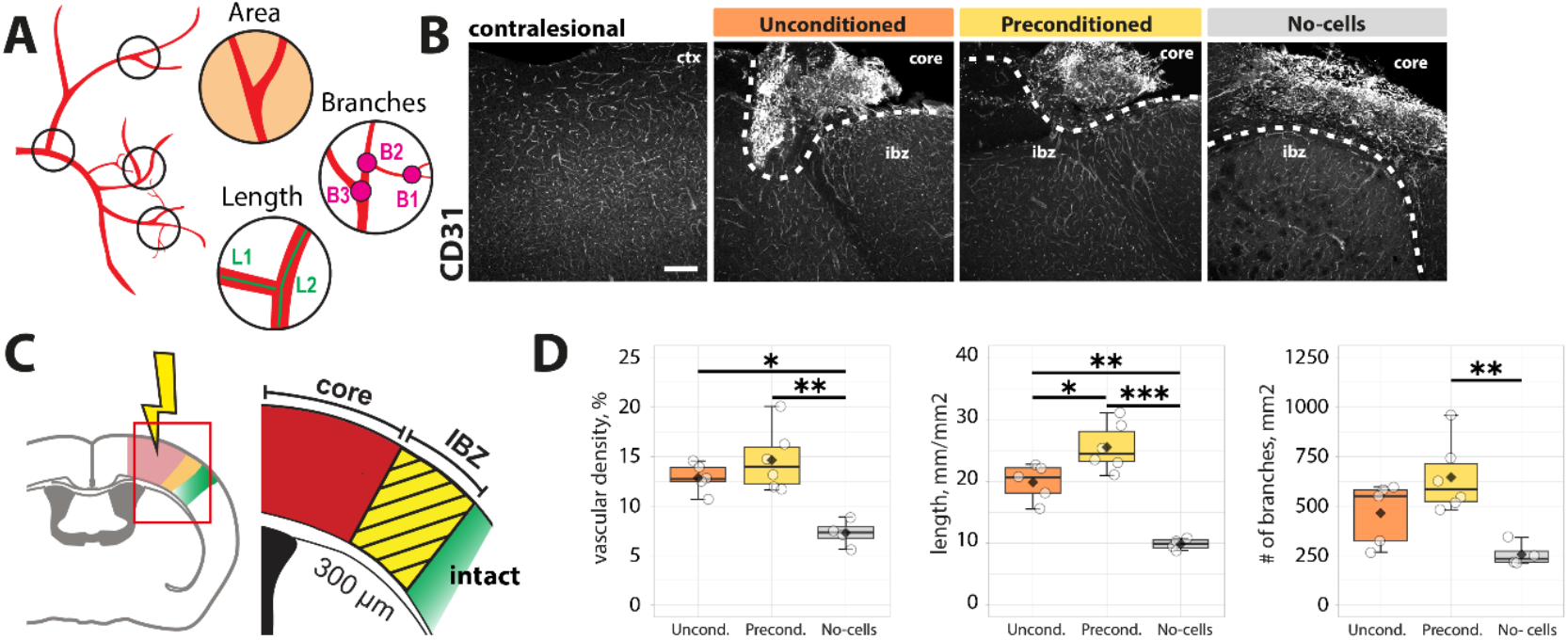
Hypoxic preconditioning of grafted cells increases angiogenesis. (**A**) Schematic defining area, length and branches in vasculature. (**B**) Representative images of vasculature stained with CD31 in the cortex (ctx) of the contralateral hemisphere (left) and of the ipsilateral site. The dotted line marks the border between the core and the ischemic border zone. Scale bar, 100 μm. (**C**) Schematic of stroke core (red) and ischemic border zone (ibz, yellow) marking a boundary to the intact tissue (green). (**D**) Quantification of vascular density (left), vessel length (middle) and number of branches (right) in the ischemic border zone. In D: Differences between unconditioned, preconditioned and no-cells control subjects were assessed using ANOVA with tukey HSD post-hoc test. Boxplots show center line = median, box = interquartile range, whiskers = minimum–maximum, and each point represents one animal. *p<0.05, **p<0.01, ***p<0.001.

These findings indicate that hypoxic preconditioning of cell grafts supports host vascular remodeling after stroke.

## DISCUSSION

In this study we show that hypoxic preconditioning of neural progenitor cell (NPC) grafts prior to transplantation represents a promising approach to improve graft survival in the ischemic brain. Hypoxia induced metabolic adaptations and altered growth and differentiation profiles *in vitro*, and locally transplanted NPCs showed increased long-term abundance after stroke. Mice receiving hypoxia-preconditioned NPCs exhibited enhanced regeneration-associated responses, including increased vascular remodeling and attenuated inflammatory signaling, compared to mice transplanted with unconditioned NPCs.

Our findings build on previous studies showing that short-term hypoxic preconditioning enhances the survival and paracrine activity of MSCs, ESCs, and NSCs after transplantation into ischemic tissue. Mechanistically, hypoxic preconditioning has been proposed to increase graft survival^7^ by activating hypoxia-inducible pathways that reduce apoptosis, increase angiogenesis, and modulate inflammation^8–10^. By applying sustained hypoxia to human iPSC-derived NPCs, we extend this concept to a clinically relevant cell source that is cultured under xeno-free and GMP-compliant conditions^13^ and demonstrate that preconditioning not only improves graft viability in the post-stroke brain but also supports long-term regeneration-associated tissue responses.

In our study, hypoxic preconditioning induced metabolic adaptations characterized by increased reliance on glycolysis and downregulation of proliferative and mitotic programs. This is in line with established cellular adaptations to hypoxia, where HIF-1α and Wnt/β-catenin signaling has been shown to promote glycolysis and NPC proliferation.^10^ During differentiation, we observed a relative increase in glial cells. Similar shifts in fate under reduced oxygen have been described previously, where hypoxia was shown to bias neural progenitors toward astrocytic lineages.^14^ Notably, we did not observe this glial shift *in vivo*, likely suggesting the influence of the complex post-stroke environment that provides additional cues including inflammatory signals directing cell graft function and fate.^15^ The predominant loss in neurons after an ischemic event in brain tissue may also account for the prevailing neural fate of cells.^16^

Cell grafts with and without hypoxic preconditioning showed similar Ki67 proliferation rates and comparable graft composition, although their total abundance differed at 4 weeks after stroke. This suggests that hypoxia-preconditioned grafts may have proliferated more actively during the early post-transplantation period, as indicated by our *in vitro* findings. Bioluminescence imaging further supported this idea since some animals transplanted with preconditioned cells showed increased signal 3 days post-injection whereas all cell grafts in control animals had a decreased cell survival. This is the most critical phase for cell survival as the cells are transferred into a hostile environment hence hypoxic preconditioning may protect cells by adapting to hypoxia. Furthermore, the hypoxia-induced glycolytic shift that we observed *in vitro*, has been previously described to enhance resistance to apoptosis in different cell types including neural stem cells^17,18^, and improved therapeutic outcome in related brain injury models.^19^ Although some studies also indicated that sustained severe hypoxia can also increase cell death in iPSC-derived cells.^20,21^

Beyond changes in grafted cells, we also found that hypoxia-preconditioned NPCs influenced the host response by promoting vascular remodeling and reducing microglial activation. These effects are consistent with previous observations by us and others that transplanted NPCs act not only through cell replacement but can also contribute to repair via trophic support and immunomodulation.^22,23^ Future studies are needed to address whether these structural and molecular changes could also lead to long-term functional recovery.

There are some limitations that need to be addressed in future studies. While our study assessed graft survival and host responses on a single post-transplantation time point of 4 weeks post-stroke, which is still considered as subacute stage of stroke, chronic conditions were not investigated. Future studies will be required to establish whether the benefits of hypoxic preconditioning persist into the chronic phase of stroke. Second, our work was performed in Rag2^-/-^ immunodeficient mouse model, which does not capture the full complexity of the human immune response after stroke. Although we previously showed that Rag2^-/-^ immunodeficient mice have comparable stroke responses to WT mice^24^, they lack adaptive immunity, and therefore potential interactions between grafted cells and host T or B cells may not be fully captured. Third, while we observed robust changes in graft survival, fate, and host responses, the mechanistic basis of these effects remains incompletely understood. Future studies are needed to dissect graft-host interactions^22,25^ to better understand the underlying pathways. In addition, the continuous expansion of cells under hypoxic conditions comes with expenditures and the effect is transient. Mimicking the effect with specific agents would simplify the workflow and ensure a stable upregulation of supportive pathways. Future studies have to be conducted to evaluate an efficient agent for consistent support.

## MATERIALS & METHODS

### Cell culture

For generating NPCs from iPSCs with a dual-reporter system, we followed the protocol previously described from our group^13^. In brief, iPSC to NPC differentiation was induced in xeno-free media containing small molecules in plates coated with Vitronectin. After 6 days, cells were replated onto poly-L-ornithine/Laminin (pLO/L521) coated plates. Neural Stem cell Maintenance Medium (NSMM, 50% DMEM/F12, 50% Neurobasal medium, 1 × N2-supplement, 1 × B27-supplement, 1 × Glutamax, 10 ng/ml hLIF, 4 µM CHIR99021, 3 µM SB431542) supplemented with 5 ng/mL FGF2 (fibroblast growth factor 2) was used for maintenance and passaging. Cells were transduced with a lentiviral vector containing bioluminescent (red firefly luciferase) and fluorescent (eGFP) reporter for *in vivo* tracking. Spontaneous neural differentiation was initiated by withdrawing small molecules and FGF2. During the experimental period NPCs were either cultured at 37°C under normoxic conditions (20% O_2_, 5% CO_2_, 75% N_2_) or in a box with a hypoxic atmosphere (1% O_2_, 5% CO_2_, 94% N_2_).

### Transcriptomics

RNA of NPCs was isolated using the RNeasy kit from Qiagen according to the manufacturer’s instructions. High throughput sequencing, including library preparation, sequencing, read processing, read alignment, and read counting, was managed and executed by the Functional Genomics Center Zurich (FGCZ), University of Zurich. Samples were delivered at a concentration of 100 ng/μL diluted in nuclease-free water. Quality control was conducted for each sample prior to sequencing. Approved samples were sequenced on an Illumina Novaseq 6000 platform using the Illumina Truseq mRNA library preparation protocol.RNA sequencing and gene set enrichment analysis were performed using EdgeR and clusterProfiler (v4.0) within the R software.

### Viability assays

To longitudinally measure cell viability during cell culture experiments, Cell Counting Kit 8 (CCK8, Abcam), which makes use of dehydrogenase activity in live cells, was applied. Cells were seeded on pLO/L521 coated 24-well plates at a density of 30’000 cells/well (day 0). The assay was performed daily during proliferation and at day 7, 11, 15, and 17 during differentiation. CCK8 solution was added directly into the media to achieve a dilution of 10% and plates were incubated for 3 h under predefined conditions. Subsequently, optical density (OD) was measured with a spectrophotometer (Infinite 200 Pro, Tecan) at 450 nm with 9 reads per well. After measuring, the solution was replaced with fresh media.

Cell viability after 18 days was determined by trypan blue exclusion method using Vi-Cell XR Viability Analyzer (Beckman Coulter). Cells were seeded on 24-well plates at 50’000 cells/well and maintained in predefined conditions before harvesting and measuring at day 18.

### Seahorse Assay

Metabolic functions in living cells were assessed using the Seahorse XFe24 Analyzer (Agilent), following the instructions of the manufacturer. 100’000 cells/well were seeded on pLO/L521 coated Seahorse culture plates and kept under normoxic conditions or in a hypoxic chamber (<5% O_2_) overnight. 1 h before starting the experiment, culture medium was replaced with Seahorse DMEM medium (103575-100, Agilent) supplemented with 10 mM glucose, 1 mM pyruvate, and 2 mM L-glutamine and the plate was incubated in a non-CO_2_ incubator. The drug solution for the assay contained 1.5 µM oligomycin and 0.5 µM rotenone + antimycin A (ROT/AA). 4 measurements at baseline and 3 measurements after drug induction were performed and the assay was repeated 3 times. Immediately after the assay, living cells were fixed with 4% PFA in PBS and stained with 20 ng/µL DAPI (4′,6-Diamidin-2-phenylindol, 1:100 in PBS, Sigma) for 10 min at room temperature (RT). Microscopic images were taken with an inverted *Zeiss* fluorescent microscope at 10x magnification. The number of nuclei was quantified with an automatized script in Fiji (ImageJ) for normalizing to cell count. Wave Desktop and Controller 2.6 Software (Version 2.6.1) was used to analyze the assay data. Data visualization was done in RStudio.

### Proliferation assay

To determine the proliferation ratio of cells, EdU (2’-Deoxy-5-ethynyluridine, Abcam) was administered to live cells. Cells were seeded on coverslips in 24-well plates at a density of 30’000 cells/well. 10 mg/mL of EdU was added directly into the media and incubated for 2 h in the proliferation phase (day 4) and 8 h for cells under differentiation (day 18) at 37°C. Cells were subsequently fixed with 4% PFA.

### Hypoxia staining assay

Image-iT™ Green Hypoxia Reagent (H-iT, Thermo Fisher) was added directly to the media to a final concentration of 5 mM. After 30 min of incubation at 37°C, the reagent was exchanged with fresh media and cells were then incubated again for 4 h under the designated oxygen condition before fixation with 4% PFA.

### Immunocytochemistry

Fixed cells were blocked for 1 h at RT with 5% donkey serum (DS) plus 0.1% Triton X-100 in 0.1 M PBS followed by overnight incubation at 4°C with primary antibodies (**Supp. Table 1**) diluted in 3% DS plus 0.1% Triton X-100 in 0.1 M PBS. Cells were washed three times with PBS prior to secondary antibody (**Supp. Table 2**) incubation for 2-3 h at RT. For EdU staining, cells were incubated for 30 min at RT with reaction buffer (ddH_2_O, 2 M Tris, 10 mM CuSO_4_, 10 mM AF 647 (Invitrogen), 500 mM ascorbic acid). Samples were counterstained with 20 ng/μl DAPI (1:100 diluted in PBS Sigma) for 10 min at RT. Cells were rewashed before mounting with Mowiol® solution on Superfrost Plus™ microscope slides. All images were captured with Leica SP8 laser scanning confocal equipped with 10X, 20X and 63X objectives. Processing of images was done in Fiji (ImageJ).

### Animals

All animal experiments were conducted according to the local guidelines for animal experimentation in the Laboratory Animal Service Center (LASC), Schlieren and were approved by the Cantonal Veterinary Office of Zurich. 9 male and 12 female (6-15 weeks, 15-30 g) immunodeficient Rag2^-/-^ knockout mice were used. Animals were housed in top-filter laboratory cages under controlled temperature and humidity conditions with a 12 h light/12 h dark cycle (lights from 6:00 a.m. to 6:00 p.m.). Mice had access to ad libitum food and water and were standard housed in groups of two to five animals per cage.

### Animal surgeries

Methods of stroke induction and cell transplantation were described previously ^24,26^. Briefly, anesthesia was induced using 4% isoflurane vaporized in oxygen. Once deep anesthesia was reached, mice were placed in a stereotactic frame (David Kopf Instruments) with a self-fabricated face mask to maintain anesthesia (1-2% isoflurane). A heating mat was placed underneath to maintain body temperature between 36-37°C. Before starting the surgery, reflexes were checked with toe and tail pinches to ensure deep anesthesia. If no reflexes were observed anymore, analgesics were injected intraperitoneally to reduce pain and to prevent inflammation after the surgery. Animals were marked on their tail for identification, and eye lubricant (Viscotears®, Bausch & Lomb) was applied to the eyes to prevent dryness. The head area between the ears and eyes was shaved with an electric razor, sanitized with Betadine (Braun), and treated with local anesthetic cream containing lidocaine and prilocaine (Emla™). Ear bars were gently inserted to fix the head in a tight position. The skull was then exposed through a midline incision, and the skin was retracted until Bregma and Lambda were visible. Bregma was located using an *Olympus SZ61* microscope and then marked as an auxiliary point. The exact location for stroke induction (AP: −1.5 to +1.5 mm and ML: 0 to +3 mm from Bregma) was determined using a stereotactic coordinate system (WPI UMP3T-1). The photosensitive dye rose Bengal 15 mg/ml dissolved in 0.9% NaCl solution was injected intraperitoneally at 10 µL/g body weight 5 min before illumination. The targeted brain region was illuminated for 10 min with a cold light source lamp (Olympus KL1500LCD, 3030 K, 150 W). Afterwards, animals were gently removed from the stereotactic frame and placed into an empty cage for post-anesthetic recovery. Stroke induction was confirmed by Laser Doppler imaging (LDI, Moors Instruments, MOORLDI2-IR). Flux data was exported and then quantified with Fiji (ImageJ) and analyzed in R software. The wound was closed using a surgical suture kit (Braun) and disinfected with Betadine®. Mice were placed in an empty cage for recovery, followed by standard post-op care.

For iPSC-NPC transplantation, cells were freshly thawed and resuspended in sterile DPBS at a concentration of 1×10^5^ cells/µL. Cell suspension was kept on ice until transplantation. Surgical procedures remained identical to those described above until locating the targeted brain area. At this stage, a small hole with a diameter of approximately 0.8 mm was drilled in the skull at identified injection coordinates (AP: + 0.8 mm / ML: + 1.5 mm (relative to bregma)). Prior to injection, the needle was gently advanced to the dorsoventral coordinates (-0.8 mm) and further moved to DV-0.9 mm for creating a pocket for the injected cells to avoid spillover. A total volume of 2 μL of the prepared cell suspension was injected at a constant injection rate of 3 nL/s using a 33 gauge needle attached to a 10 μL Hamilton syringe. Following injection, the needle was left in place for another 5 min before a slow retraction to prevent reflux and loss of cells. Afterwards, Histoacryl® was applied to seal the burr hole, and the wound was sutured. Mice were returned to a cage with a warming pad and received standard post-operative care.

### Bioluminescent imaging (BLI)

*In vivo* Bioluminescence imaging (BLI) was carried out to longitudinally track the injected cells. Transplanted animals were injected intraperitoneally with 300 mg/kg body weight D-Luciferin dissolved in NaCl (0.9%, Braun) 5 min before isoflurane anesthesia (4% induction, 1.5%-2.5% maintenance). Images were acquired with the IVIS imaging system (Perkin Elmer) at a 5 min interval starting 10 min after substrate injection using automated settings. BLI data were analyzed using the Living Image software (V 4.7.3) by measuring the radiance in a region of interest (ROI) that covered 1.8 cm^2^ over the head. Background BLI signal from a randomly selected region on the animal’s back was subtracted from all the values. Data was visualized with R software.

### Immunohistochemistry

For immunohistochemistry, animals were perfused with 100ml Ringer solution followed by 100ml 4% PFA solution. Brain tissue was removed from the skull cavity and treated with 4%PFA for 6 h before incubating in 30% sucrose. Fixed brains were cut into coronal slices with a thickness of 40 µm using a microtome (Thermo Scientific HM 450). Brain slices were washed with 0.1 M PBS before incubation with blocking buffer containing 5% donkey serum diluted in 0.1 M PBS + 0.1% Triton® X-100 for 1 h at RT. Subsequently, primary antibodies (**Supp. Table 1**) were added and incubated at 4°C overnight on a shaker. The next day, sections were washed with 0.1 M PBS and then incubated with the corresponding secondary antibodies (**Supp. Table 2**) for 2 h at RT. DAPI (20ng/μL, 1:100 diluted in PBS, Sigma) was used for counterstaining of cells. Sections were mounted onto Superfrost Plus™ microscope slides with Mowiol® mounting solution. Images were acquired with confocal microscopy (Leica SP8 confocal) and processed in Fiji (ImageJ).

### Microscopy of stroke area/volume and graft volume quantification

Imaging of brain sections was carried out on a Leica SP8 laser-scanning confocal microscope using 10X and 63X objectives, and images were processed in Fiji (ImageJ) as described previously ^27^. For stroke and graft size quantification, 40 μm coronal sections, stained for HuNu (graft) and GFAP (lesion), were scanned on a Zeiss Axio Scan.Z1 slide scanner and subsequently analyzed in Fiji. Stroke size was assessedby manually tracing a polygon tightly around the infarct to delineate the lesion area per section. For graft size analysis, cell distribution was delineated in two distinct areas: the “graft center”, defined as the region with densely populated cells (>3x HuNu signal to background), and the “graft periphery”, where cell density was reduced (HuNu signal threshold between > 1.5x and < 3x to background). Sections were assigned to stereotaxic levels (distance relative to bregma) using a mouse brain atlas^28^. A custom R script was used to rescale images and to enhance contrast. Lengths, widths, and areas were measured on each section and combined to generate a 3D representation to estimate lesion and graft volumes. These volumes were then approximated as elliptical oblique cones (1):

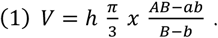

### Histological quantification of vasculature, microglia and astrocytes

Post-ischemic angiogenesis was analyzed in the peri-infarct region (ischemic border zone, IBZ) adjacent to the stroke core, extending up to 300 μm. Quantification was done in Fiji (ImageJ) using a previously validated script to automatically compute (a) vascular density, (b) branch and junction counts, and (c) vessel length ^29–31^. Glial scar intensity and inflammatory signal were assessed on three sections per animal immunostained for GFAP and Iba1. Images were converted to 8-bit and thresholded using mean gray values from regions of interest in the unaffected contralateral cortex to generate binary masks. Then, the cumulative area of reactive gliosis and inflammation within and around the ischemic lesion was calculated.

### Statistical analysis

Statistical analysis was carried out in RStudio (Version 4.3.2). All data was tested for normal distribution using Shapiro-Wilk test. An unpaired two-sided t-test was applied to compare normally distributed data with two independent groups. Data of cerebral perfusion did not follow a normal distribution and therefore tested with a paired Wilcoxon test. ANOVA was used to for multiple comparisons. The significance between means of groups in multiple comparisons was identified using TukeyHSD post-hoc test. Estimated marginal means (EMM) post-hoc test with Bonferroni correction was carried out to compare bioluminescence data of animals over the timecourse. Significance was defined as *p < 0.05, **p < 0.01, ***p < 0.001, ****p <0.0001.

## Acknowledgements

This work is supported by the funding from Swiss 3R Competence Center (OC-2020-002), the Swiss National Science Foundation (CRSK-3_195902 and PZ00P3_216225), the Neuroscience Center Zurich, USC Dean’s Pilot Award, and the Mäxi Foundation.

## Author contributions statement

NHR, RZW, CT, and RR contributed to the overall project design. NHR, RZW, RR, CB, and KJZ conducted and analyzed in vitro experiments. VB, NHR performed energy metabolism assays. NHR, RZW, and RR conducted and analyzed in vivo experiments. RR analyzed RNAseq data. NHR, RZW, CT, and RR prepared figures. MG provided an iPS cell line. CT and RR supervised the study. NHR, RZW, CT, and RR wrote and edited the manuscript with input from all authors. All authors read and approved the final manuscript.

