## Supplementary material for "Hypoxia preconditioned neural xenografts promote repair of brain tissue after stroke": Supp Tables

**Suppl. Table 1: Primary antibodies used for immunofluorescence stainings.**

| **Antibody** | **Material Nr.** | **Specificity** | **Host** | **Supplier** | **Dilution** |
| --- | --- | --- | --- | --- | --- |
| Oct4 | 701756 | Stem cell marker | Rabbit | Invitrogen | 1:300 |
| Nestin | MAB1259 | Marker for human neural progenitor cells | Mouse | R&D Systems | 1:150 |
| GFAP | BP5082 | Binds to Glial fibrillary acidic protein (GFAP)in astrocytic cells. | Guinea pig | Origene | 1:100 |
| β-III-Tubulin | MAB5568 | Recognizes βIII-Tubulin in neural cells. | Rabbit | Cell Signaling | 1:200 |
| HuNu | MAB1281 | Detects human nuclei | Mouse | Sigma-Aldrich | 1:200 |
| Nanog | AF1997 | Pluripotency marker | Goat | R&D Systems | 1:30 (ICC), 1:100 (IHC) |
| Pax6 | ab5790 | Marker for human neural progenitor cells | Rabbit | Abcam | 1:200 |
| Ki67 | MA5-14520 | Detects proliferating cells | Rabbit | Invitrogen | 1:150 |
| GFAP | EPR1034Y | Binds to Glial fibrillary acidic protein (GFAP)in astrocytic cells. | Mouse | Abcam | 1:500 |
| CD31 | 550274 | Marker for blood vessels | Rat | BD Biosciences | 1:50 |
| Iba1 | ab5076 | Microglia marker | Goat | Abcam | 1:200 |
| βIII-Tubulin | PA5-85639 | Marker for axons of neurons | Rabbit | Invitrogen | 1:200 |
| Synaptophysin | MA5-14532 | Presynaptic vesicles | Rabbit | Invitrogen | 1:200 |

**Suppl. Table 2: Secondary Antibodies used for immunofluorescence stainings.**

| **Antibody** | **Material Nr.** | **Fluorophore** | **Host** | **Supplier** | **Dilution** |
| --- | --- | --- | --- | --- | --- |
| Anti-mouse IgG | 715-545-151 | Alexa Fluor 488 | Donkey | Jackson | 1:500 |
| Anti-rabbit IgG | 711-165-152 | Cy3 | Donkey | Jackson | 1:500 |
| Anti guinea pig IgG | 706-165-148 | Cy3 | Donkey | Jackson | 1:500 |
| Anti-goat IgG | 705-165-147 | CY3 | Donkey | Jackson | 1:250 |
| Anti-mouse IgG | 715-165-150 | CY3 | Donkey | Jackson | 1:500 |
| Anti-rabbit IgG | 711-605-152 | Alexa Fluor 647 | Donkey | Jackson | 1:100 |
