## Supplementary material for "Hypoxia preconditioned neural xenografts promote repair of brain tissue after stroke": Supp Figures

### CORRESPONDANCE

**Ruslan Rust, Ph.D.**
Assistant Professor
The Zilkha Neurogenetic Institute,
Department of Physiology and Neuroscience
Keck School of Medicine of the University of Southern California
1501 San Pablo Street
Los Angeles, CA 90033

ORCID: 0000-0003-3376-3453

**Christian Tackenberg, Ph.D.**
Scientific Head of Division Neurodegeneration
Institute for Regenerative Medicine • IREM
University of Zurich
Wagistrasse 12
8952 Schlieren, Switzerland

ORCID: 0000-0002-0019-3055

A

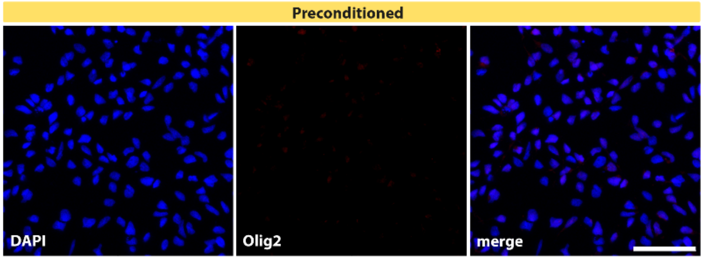


**Suppl. Figure 1: Phenotyping of NPCs after 14 days of neural induction.** (**A**) Immunofluorescence for Olig2 (not detected) counterstained with DAPI (blue) in preconditioned cells. Scale bar, 50 µm.

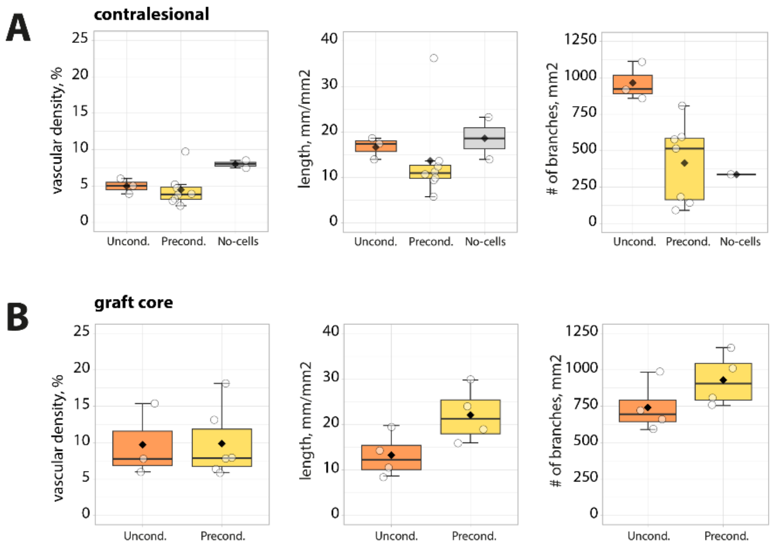


**Suppl. Figure 2: Quantification of angiogenesis in contralesional site and graft core.** (**A,B**) Quantification of vascular density (left), vessel length (middle) and number of branches (right) in the contralesional site and in the graft core. Differences between unconditioned, preconditioned and no-cells control subjects were assessed using ANOVA with tukey HSD post-hoc test. Boxplots show center line = median, box = interquartile range, whiskers = minimum–maximum, and each point represents one animal.
